# Spontaneous eye blink rate predicts individual differences in exploration and exploitation during reinforcement learning

**DOI:** 10.1101/661553

**Authors:** Joanne C. Van Slooten, Sara Jahfari, Jan Theeuwesu

## Abstract

Spontaneous eye blink rate (sEBR) has been linked to striatal dopamine function and to how individuals make value-based choices after a period of reinforcement learning (RL). While sEBR is thought to reflect how individuals learn from the negative outcomes of their choices, this idea has not been tested explicitly. This study assessed how individual differences in sEBR relate to learning by focusing on the cognitive processes that drive RL. Using Bayesian latent mixture modelling to quantify the mapping between RL behaviour and its underlying cognitive processes, we were able to differentiate low and high sEBR individuals at the level of these cognitive processes. Further inspection of these cognitive processes indicated that sEBR uniquely indexed explore-exploit tendencies during RL: lower sEBR predicted exploitative choices for high valued options, whereas higher sEBR predicted exploration of lower value options. This relationship was additionally supported by a network analysis where, notably, no link was observed between sEBR and how individuals learned from negative outcomes. Our findings challenge the notion that sEBR predicts learning from negative outcomes during RL, and suggest that sEBR predicts individual explore-exploit tendencies. These then influence value sensitivity during choices to support successful performance when facing uncertain reward.

## Introduction

During our life we learn a lot by trial and error. When cooking a new dish, we learn from the feedback we receive about the outcome and change our future actions by repeating those dishes that tasted good. How we learn from interacting with our environment can be captured by reinforcement learning (RL) theory, which describes the mapping of situations to actions in order to maximise reward^1^. The neuromodulator dopamine (DA) plays an important role in how individuals learn from their interactions with the environment^2,3^ and has also been linked to individual variability in spontaneous eye blink rate (sEBR)^4-6^. While research suggest that sEBR reflects the extent to which individuals learn from negative outcomes of their actions ^5^, this idea has not been tested explicitly. Here, we set out to address this issue by associating sEBR to individual differences in how we exploit actions that likely produce desirable outcomes and learn from positive and negative feedback: the cognitive mechanisms that drive RL.

More than 30 years of research has shown that sEBR, or the frequency of blinks per unit time, is affected by DA, particularly in the striatum (for a recent review, see^7^). In general, pharmacological studies in animals and humans have shown that DA-enhancing drugs elevate sEBR, while DA-decreasing drugs suppress them ^4,6,8-12^. Moreover, sEBR is altered in clinical conditions that are associated with dysfunctions of the DAergic system ^13,14^. For example, sEBR is decreased in Parkinson’s disease (PD)^15,16^ a condition characterised by depleted striatal DA levels. These findings align with animal studies showing that MPTP - a DAergic neurotoxin that induces Parkinsonian symptoms - reduced blink rates^17^ in proportion to the post-mortem measured DA concentrations in the caudate nucleus^18^. Together, these studies generally indicate that sEBR is positively related to striatal DA function. As sEBR is a non-invasive, easily accessible measure, it can be used as a reliable yet non-specific marker of DA function. Still, it remains to be determined to which specific aspects or functions of the DA system sEBR relates^19,20^.

Recent studies have touched upon how sEBR, as a behavioural measure of individual differences in striatal DA function, relates to learning by observing links with punishment^5,6^ and reversal learning^21^. In particular, two studies found that sEBR predicted RL effects on future value-based choices^5,6^. In one of these, Slagter et al. (2015) employed a probabilistic RL task consisting of a learning and test phase. During learning, participants learned the value of different options using probabilistic feedback. Value learning was tested in a subsequent test phase where participants’ ability to avoid the least rewarded option and to approach the most rewarded option was evaluated. They found that individuals with a lower sEBR were better at avoiding the least rewarded option, while individuals with a higher sEBR were not better at approaching the most rewarded one. Thus, sEBR correlated negatively with the extent to which participants avoided the least rewarded option. The authors concluded that sEBR predicted learning from negative, but not positive, outcomes during earlier RL. However, the relation between sEBR and earlier RL was not explicitly studied, as only choices from the test phase were evaluated, and at that stage, learning had already been internalised.

Formal learning theories posit that different cognitive processes contribute to learning^1^: the learning rate determines the magnitude by which individuals update their beliefs about the environment after positive or negative outcomes, and their explore-exploit tendency describes the sensitivity to exploit actions that likely result in reward. But these different processes can have similar effects on final learned behaviour: avoiding the least-rewarded option in the test phase could be caused by both enhanced learning from negative outcomes (negative learning rate) and an exploitative choice strategy focused at avoiding negative outcomes (explore-exploit tendency). This makes previous findings^5^ ambiguous regarding which specific cognitive processes sEBR reflects.

Extending the work of Slagter et al. (2015), the current study sought to understand how sEBR relates to learning by focussing on the underlying cognitive processes that drive learning **(Figure la).** To specify these underlying processes, we used a hierarchical Bayesian version of the Q-learning RL model^22-24^ **(Supplementary Figure 1a).** This model separates RL into two different functions: an update function that updates the value of options by learning from reinforcement and a choice function that uses those learned values to guide decisions between differently valued options. The choice function calculates the probability of choosing one option over the other (e.g. option A over B), based on an individual’s sensitivity to the value difference of presented options, or explore-exploit tendency (*β*; **Figure 1b**). The outcome function updates value beliefs by reward prediction errors, which reflect the difference between predicted and actual rewards. The degree to which reward prediction errors update value beliefs is scaled by the learning rate^25^ (*α*; **Figure 1b**). As value beliefs are differently updated after positive and negative reinforcement via striatal D1 and D2 receptors ^26^ we defined separate learning rate parameters for positive (*α*_*Gain*_) and negative (*α*_*Loss*_) feedback ^23,24,27-30^.

**Figure 1:**
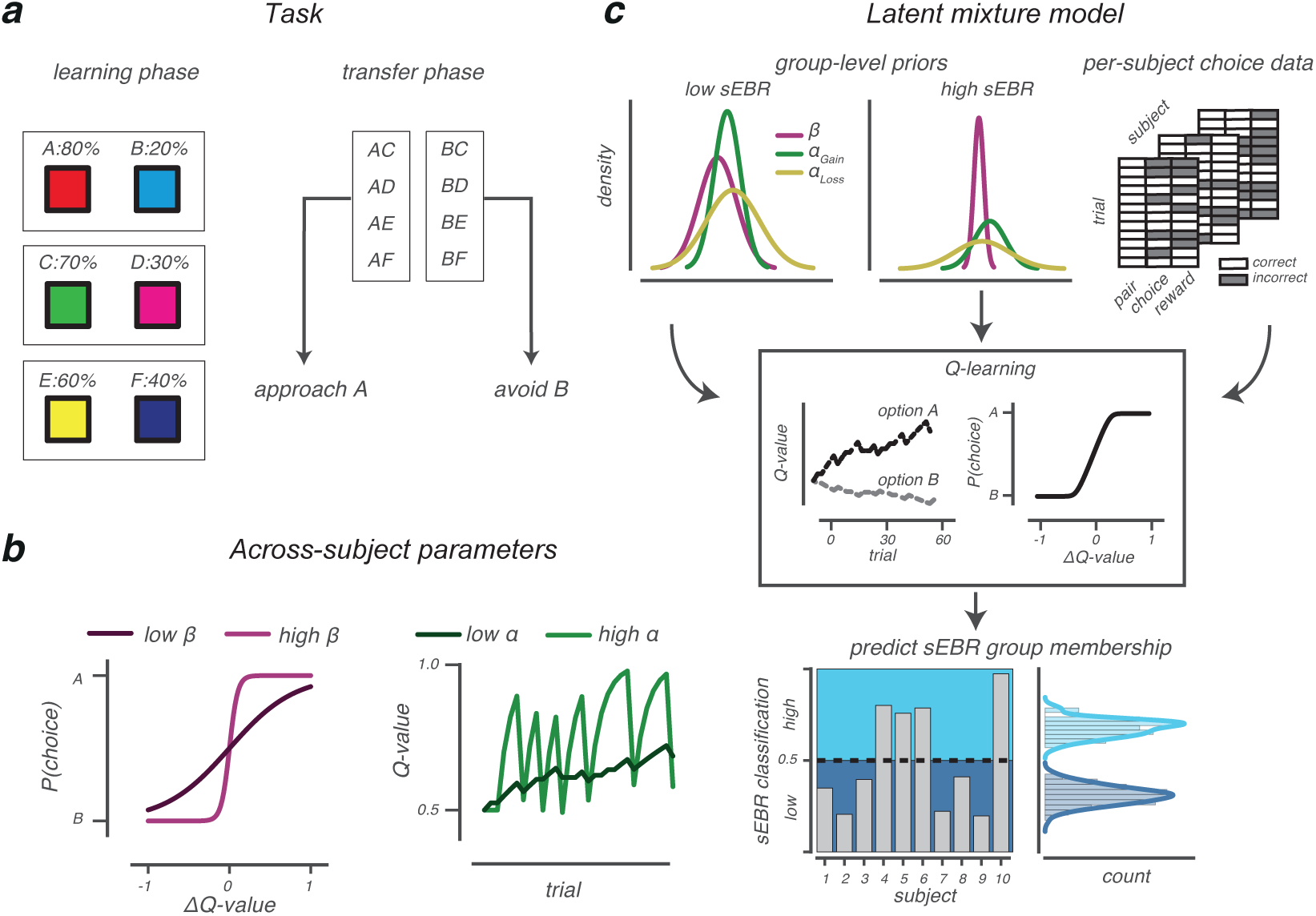
Task and model. **(a):** In the learning phase (*left*), three different option pairs (AB, CD and EF) were presented in random order and participants had to learn to select the most rewarding option of each pair (A, C and E). Each choice was followed by probabilistic auditory feedback indicating they earned a reward (+0.1 points) or no reward (no points). The probability of receiving a reward is presented for each option. The transfer phase (*right*) tested how value-based choices were influenced by earlier learning. All options were randomly paired with one another, and participants selected the most rewarding option based on previous learning, importantly, while choice feedback was omitted. The ability to approach the most rewarding option A and to avoid the least rewarding option B was evaluated, as the latter behaviour has been linked to sEBR^5^. **(b):** The *β*-parameter (*left*) describes how one’s sensitivity to option value differences (ΔQ-value) influences value-based choices. High *β*-values indicate more sensitivity to ΔQ-value, hence, more exploitatory choices for high reward options. The learning rate (*α*-parameter; *right*) describes how beliefs are updated after feedback. High learning rates indicate rapid but also volatile belief updating compared to lower learning rates. Note that only one learning rate is depicted for simplicity. **(c):** Cartoon of our Bayesian latent mixture model analysis, which we used to assess whether a participant’s sEBR (low or high) could be predicted on the basis of the estimated cognitive processes (*α*_*Gain*_, *α*_*Loss*_ and *β*) that described learning. Group-level priors were obtained from fitting a hierarchical Bayesian Q-learning model separately for low and high sEBR groups. Subsequently, the group-level priors and choice data from all participants were used as input to the latent mixture model where, critically, sEBR group membership was left out. The latent mixture model estimated for each participant the cognitive processes that described learning (using Q-learning) and calculated the probability that this participant belonged to either low or high sEBR group, given observed learning.

To our knowledge, this is the first study that directly assesses how sEBR relates to individual differences in learning. Using Bayesian latent mixture modelling techniques ^31^ **(Figure 1c** and *Methods)*, we quantify the cognitive processes that underlie learning and show that individuals with high and low sEBR can be distinguished on the basis of these cognitive processes. We then evaluate how variability in each underlying cognitive process uniquely relates to individual differences in sEBR, thereby controlling for the effects of all other variables with a network approach. We find that sEBR uniquely reflects an individual’s explore-exploit tendency (*β*), but not the tendency to learn from negative feedback (*α*_*Loss*_)- These results suggest that sEBR can be used as an easy to measure behavioural index of an individual’s explore-exploit tendency, that in turn affects the sensitivity to value differences at the time of a value-based choice.

## Results

### Blinking

On average, participants blinked 12 times per minute (median=l0.6; SO=8.3, range=l.3-34.9; **Figure 2a)**, a rate that is comparable to earlier reports^5,32,33^. When dividing participants into low and high sEBR groups based on a median split of across-subject sEBR values, low sEBR individuals blinked 5.8 times per minute (SO=2.7, range = 1.3-9.3), whereas high sEBR individuals blinked 18.3 times per minute (SO=7.3, range = 11.9-34.9). Females blinked numerically more than males (13 times versus 9 times per minute), however, their sEBR did not significantly differ (*t*(19.8)=1.26, *P=*.22, Welch’s *t*-test; *BF*_10_=0.61).

**Figure 2:**
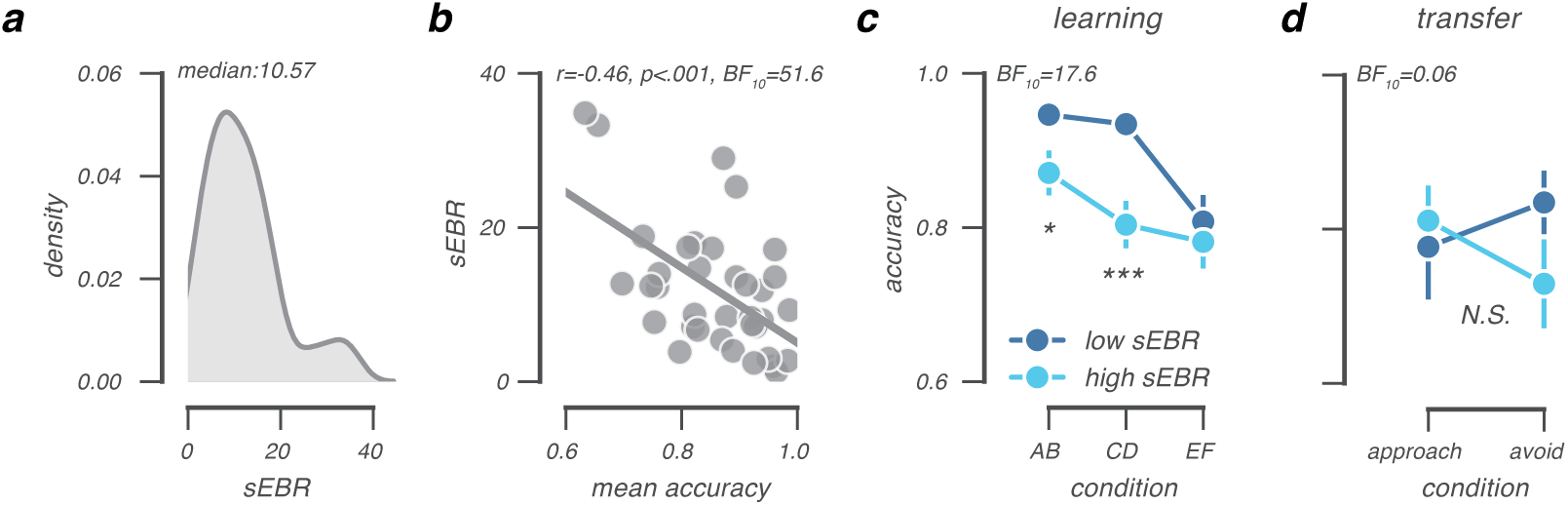
sEBR data and choice performance in the learning and transfer phase. **(a):** sEBR distribution across participants (N=36), recorded prior to the probabilistic RL task. **(b):** Lower sEBR predicts better overall choice accuracy in the learning phase. This correlation was explained by higher choice accuracy in the AB and CD pairs, but not in the EF pair **(c). (d):** In the transfer phase, choice performance was visually comparable to previous research ^5^, but there was no reliable difference benveen low and high sEBR groups in how they approached the most rewarded option and avoided the least rewarded option. *P*<.*05;* **P<.001; BF10=evidence in favour of the alternative model.

### Behavioural differences between low and high sEBR groups

Participants with low and high sEBR performed differently in the learning phase of the probabilistic RL task. Overall, lower sEBR predicted better learning phase performance (*r*=-0.46, *P=.005; BF*_10_=12.52, **Figure 2b**). As shown in **Figure 2c**, this difference was further evidenced by a mixed AN OVA with factors accuracy (AB, CD, EF) and sEBR which again showed better overall learning performance at lower sEBR (*F*(l,34)=7.23, *P*=.01; *BF*_10_=17.6), and a trend towards an interaction effect (*F*(2,68)=2.66, *P*=.08). This indicated that lower sEBR related to better learning performance in the more certain AB (*t*(34)=-2.5, *P*=.02; *BF*_10_=3.18) and CD pairs (*t*(34)=-3.7, *P*<.001; *BF*_10_=39.4), but not in the uncertain EF pair (*t*(34)=-0.5).

In the transfer phase, participants were able to approach the most rewarded option (approach-A: mean accuracy=80%, SD=24%) and to avoid least rewarded option (avoid-B: mean accuracy=79%, SD=21%) well above chance (one-sample *t-test;* both P-values<.001), indicating they successfully used previously learned option values in novel choice contexts. Overall, participants were equally successful at approach-A and avoid-B choices (*F*(l,35)=0.05). Nevertheless, we observed a pattern that numerically replicated Slagter et al. (2015), such that lower sEBR related to better avoid-B performance. Importantly, however, we did not find enough evidence for a reliable effect within this sample, as the observed interaction did not reach significance (*F*(1,34)=1.79, *P*=0.2; *BF*=5.5 in favour of the null-model, **Figure 2d).**

As fatigue is tied to poorer task performance and increased blink rates and blink durations ^34,35^ we addressed the possibility that differences in fatigue explained why individuals with a higher sEBR performed worse on the learning task. To exclude this possibility, we examined how participants’ median blink durations related to learning phase choice accuracy and sEBR. If fatigue affected choice performance, median blink durations should negatively predict learning phase choice accuracy and positively predict sEBR. Neither of these relationships were observed, as median blink durations did not correlate with learning phase choice accuracy (*r*=0.11, *P*=.51), nor with sEBR (*r*=0.15, *P*=.35), indicating that sEBR indexes individual differences in probabilistic learning that cannot be explained by fatigue.

To summarise, our behavioural results suggest that individual variability in sEBR relates to how participants learn from probabilistic feedback, with lower sEBR predicting better learning, especially from more reliable feedback.

### Q-learning parameter estimation for low and high sEBR groups

Our behavioural analysis suggested that variability in sEBR relates to how individuals learn from probabilistic feedback. To understand how this relationship is associated with, or shaped by, the cognitive processes that drive learning, we analysed choices in the learning phase of low and high sEBR groups using a Bayesian hierarchical Q-learning model **(Supplementary Figure 1**).

As shown in **Figure 3**, we observed shifts between the high and low sEBR groups in the group-level posterior distributions of all parameters, but particularly for the *β*- and *α*_*Loss*_-parameter. These observations suggested that the low sEBR group exploited high value options more often (higher-parameter) and updated value beliefs stronger after negative feedback (higher *α*_*Loss*_-parameter). Note, however, that these observations were based on visual inspections of the group-level posteriors. To formally test whether high and low sEBR groups can be distinguished on the basis of the observed differences in the estimated Q-learning parameters, we used a recently developed Bayesian latent mixture modelling approach^36^ that we adapted for Q-learning **(Figure 1c).**

**Figure 3:**
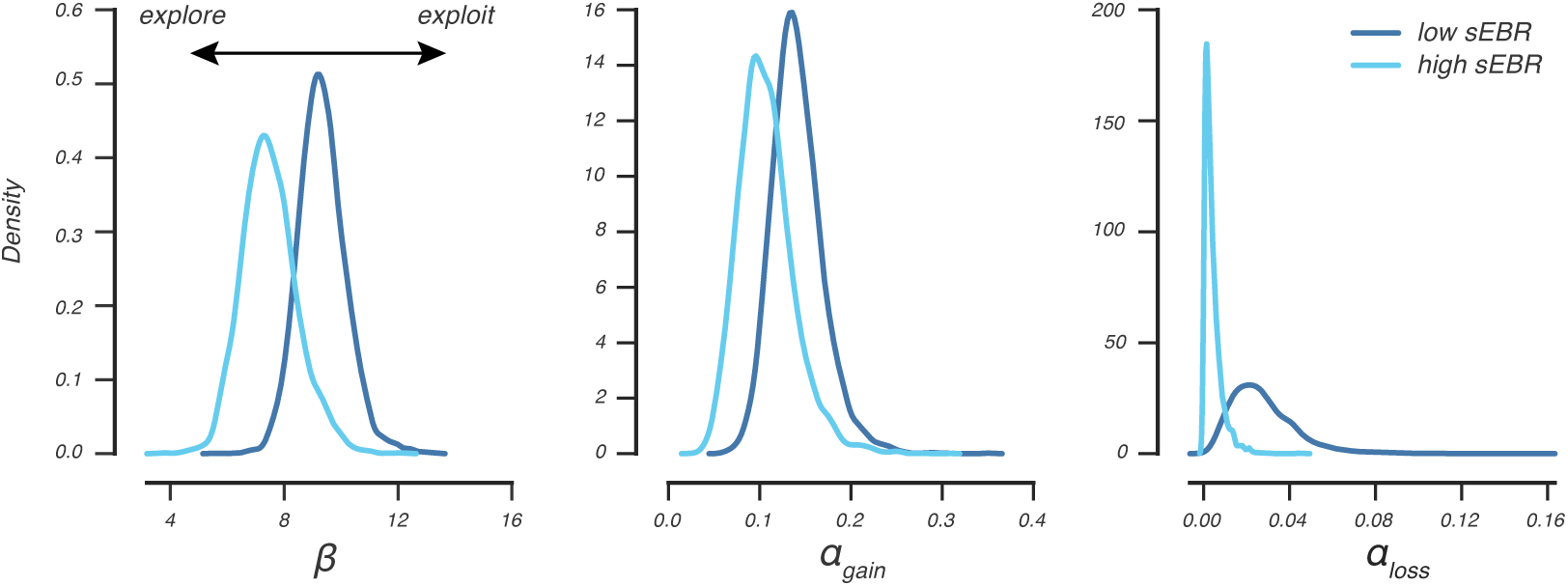
Q-learning parameter estimation for low and high sEBR groups. Posterior distributions of group-level parameters for high and low sEBR groups obtained by fitting the Bayesian hierarchical Q-learning model separately for both groups.

### Classifying sEBR group membership using Bayesian latent mixture modelling

To test whether an individual’s sEBR group membership (i.e. low or high) could be predicted solely on the basis of the estimated Q-learning parameters (*α*_*Gain*_, *α*_*Loss*_ and *β*), we implemented a two-group Bayesian latent mixture model **(Fig 1c** and *Methods* for a detailed description of this approach).

As shown in **Figure 4a**, our Bayesian latent mixture model correctly classified 72% of participants using the estimated Q-learning parameters, a percentage that was well above chance (*P*=.011, *BF*_10_=14.5; *one-sided binomial test*). Consistently, higher probabilities to be classified as a member of the high sEBR group by the latent mixture model predicted higher sEBR values (*r*=0.51, *P*<.001, *BF*_10_=24.9, **Figure 4b**), which effectively shows that the learning-based mixture classification positively related to sEBR measurements that were recorded prior to the probabilistic RL task. Together, these results highlight that low and high sEBR groups can be distinguished on the basis of the cognitive processes they relied on during learning.

**Figure 4:**
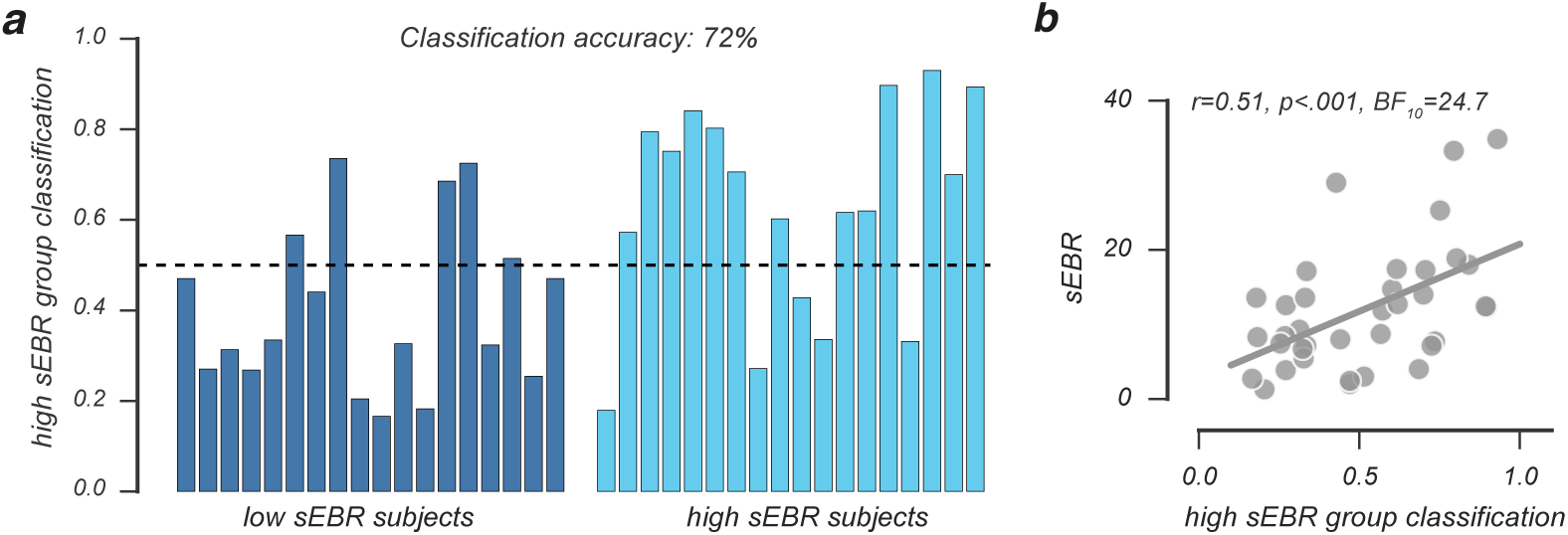
Bayesian latent mixture model classification of sEBR group membership. **(a)** Per-participant posterior classification probability to belong to the high sEBR group. A low posterior classification probability suggest that a participant is very likely to fall into the low sEBR group, whereas a high posterior classification probability indicates the participant very likely belongs to the high sEBR group. **(b)** The probability to be classified into the high sEBR group correlated positively with sEBR measurements.

### sEBR predicts individual differences in exploration and exploitation

Our prior analyses showed that sEBR relates to differences in learning that were driven by a differential use of underlying cognitive processes. However, it remains unknown what the relative influence is of each cognitive process on sEBR, leaving open the question how sEBR relates to individual variability in how we update our beliefs after desired (*α*_*Gain*_) and undesired (*α*_*Loss*_) outcomes, or the variability by which we exploit actions that will likely result in reward (*β*).

We used a multiple regression model that incorporated all three cognitive processes (*α*_*Gain*_, *α*_*Loss*_ and *β*) to explain individual variability in sEBR. The model well accounted for the variability in sEBR (*F*_(3,32)_ = 5.8, *P* = .003, *R*^2^ = 0.35), which was driven by a significant contribution of the *β*-parameter (*b*_*β*_ (SE)=-4.5 (1.2), *z*=-3.7, P<.001, *BF*=33.8), but not the 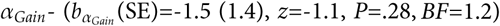 or the *α*_*Loss*_-parameter 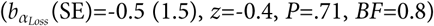. As shown in **Figure 5a**, the Bayesian linear regression analysis further indicated that the model that only incorporated the *β-* parameter to explain individual variability in the sEBR data was 47 times more likely to explain the data compared to the null-model, which is regarded very strong evidence in favour of this model^37^ **(Supplementary Table 1). Figure 5b** illustrates the negative relationship between the *β*-parameter and sEBR, indicating that exploitative decision makers had a lower sEBR. Together, these results link sEBR to individual variability in exploiting actions that lead to rewarding outcomes, but not to the magnitude by which individuals update their value beliefs after positive or negative outcomes.

**Figure 5:**
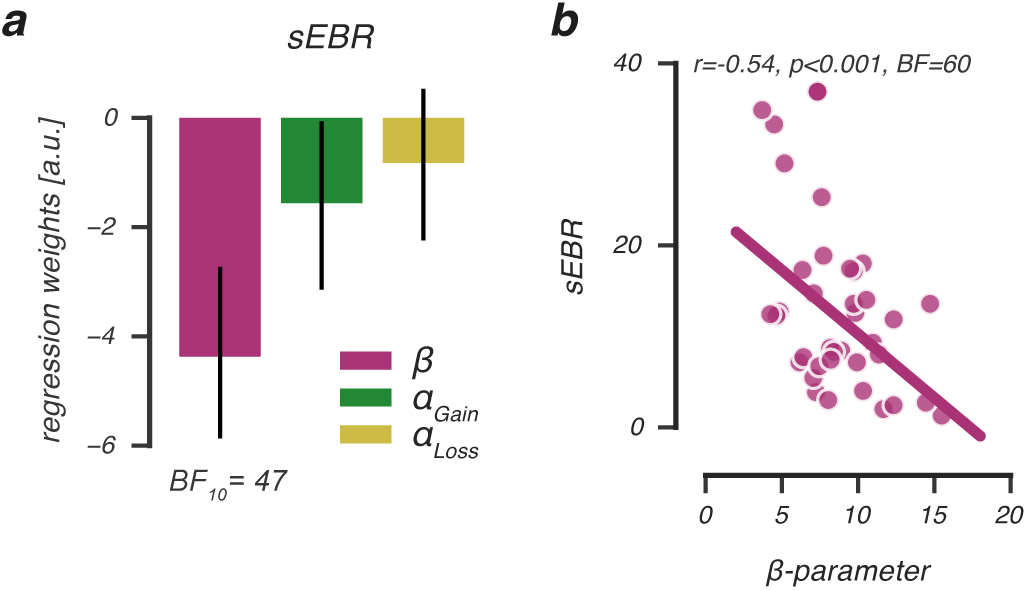
sEBR predicts individual differences in exploration and exploitation. **(a)** Beta coefficients of a multiple regression analysis indicating that *β*-parameter estimates uniquely and negatively relate to sEBR. This was further illustrated by a negative correlation between individual *β*-parameter estimates and sEBR **(b)**, showing that low sEBR individuals exploited highly values options more often compared to high sEBR individuals.

### Learning effects on choices in the transfer phase

Our results thus far relate sEBR to how participants make value-based choices during learning, but show no reliable effect of sEBR on avoid-B or approach-A choices in the transfer phase. Because this relationship has been reported in the past^5^ this section additionally examined how the Q-learning parameters (*α*_*Gain*_, *α*_*Loss*_, *β*), related to approach-A and avoid-B performance in the transfer phase.

Results from a multiple regression analysis indicated that individual variability in avoid-B, but not approach-A, performance was predicted by the Q-learning model parameters *F*_(3.32)_ = 3.7, *P* = .02, *R*^2^ = 0.26; **Figure 6**). This was driven by a significant contribution of the *β*-parameter (*b*_*β*_(SE)=0.069 (0.03), *z*=2.066, *P*=.047, *BF*=2.4) and a smaller, albeit non-significant, contribution of the *α*_*Loss*_-parameter 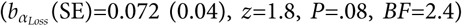. The Bayesian linear regression analysis further indicated that a model that incorprated both the *β*- and *α*_*Loss*_-parameter as main factors to explain individual variability to avoid-B performance was 7 times more likely to explain the data compared to the null-model, and 3.7 times more likely compared to all other candidate models **(Supplementary Table 2)**. Together, these analyses show that an exploitative decision-making style and enhanced updating after negative outcomes predicts better avoid-B performance in the transfer phase. These results suggest that the ability to avoid undesirable outcomes is related to how individuals learn, but is unrelated to their sEBR.

**Figure 6:**
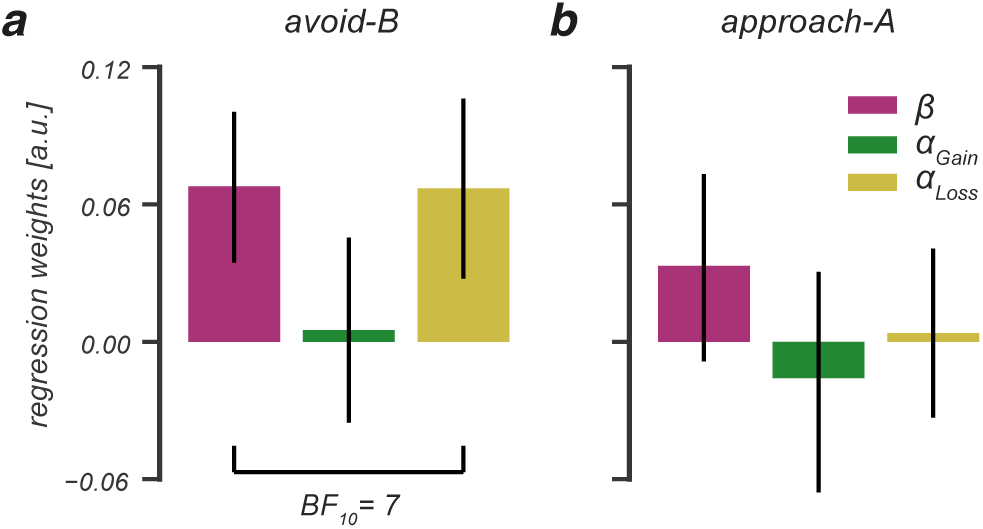
Avoid-B, but not approach-A choices in the transfer phase, are related to individual variability in negative learning rates and explore-exploit tendencies during learning. **(a)** Exploitation of high valued options (high *β*) and enhanced learning from negative feedback (high *α*_*Loss*_) during learning related to better performance to avoid the least rewarded option in the transfer phase. **(b)** Approaching the most rewarded option was unrelated to the cognitive processes that underlie learning.

### Network interactions between sEBR, cognitive learning processes and choices in the transfer phase

sEBR uniquely predicted an individual’s tendency to exploit high valued options during learning, but not approach-A or avoid-B performance given prior learning. However, individual differences in avoid-B performance were associated both with *β* (which also predicted sEBR during learning) and *α*_*Loss*_ (which is hypothesized to be associated with variability in sEBR^5^). To understand the association between these variables across learning and transfer phases, we assessed all relationships directly in one model using a network analysis.

In this final analysis, each connection in the network represents a partial correlation coefficient between two variables after conditioning on all other variables in the network. Thus, each coefficient encoded the unique association between two nodes after controlling for all other information possible^38^. **Supplementary Table** 3 shows all partial correlations between the variables, which are graphically depicted in **Figure 7.** In this graph, three important between-node relationships were observed. First, individual differences in sEBR were significantly and negatively related to the *β*-parameter (*partial r*=-0.515, *P*<.001), consistent with our previous finding that exploitative decision makers had a lower sEBR. Second, the *α*_*Gain*_ - and *α*_*Loss*_-parameter were significantly and positively related to each other (*partial r*=0.522, *P*<.001), but not to sEBR, which is inconsistent with earlier work that hypothesized sEBR indexes how much individuals learned from the negative outcomes of their choices^5^. Lastly, the ability to avoid the least rewarded option in the transfer phase related to the *β* - and *α*_*Loss*_-parameter, consistent with our previous results. However, the network analysis indicated these relationships were not robust. More importantly, the ability to avoid the least rewarded option was unrelated to sEBR, an observation that is not in line with earlier work^5^. Overall, this analysis paints a clear picture of how sEBR relates to learning and subsequent value-based choices, namely that it uniquely reflects a decision maker’s explore-exploit tendency during learning.

**Figure 7:**
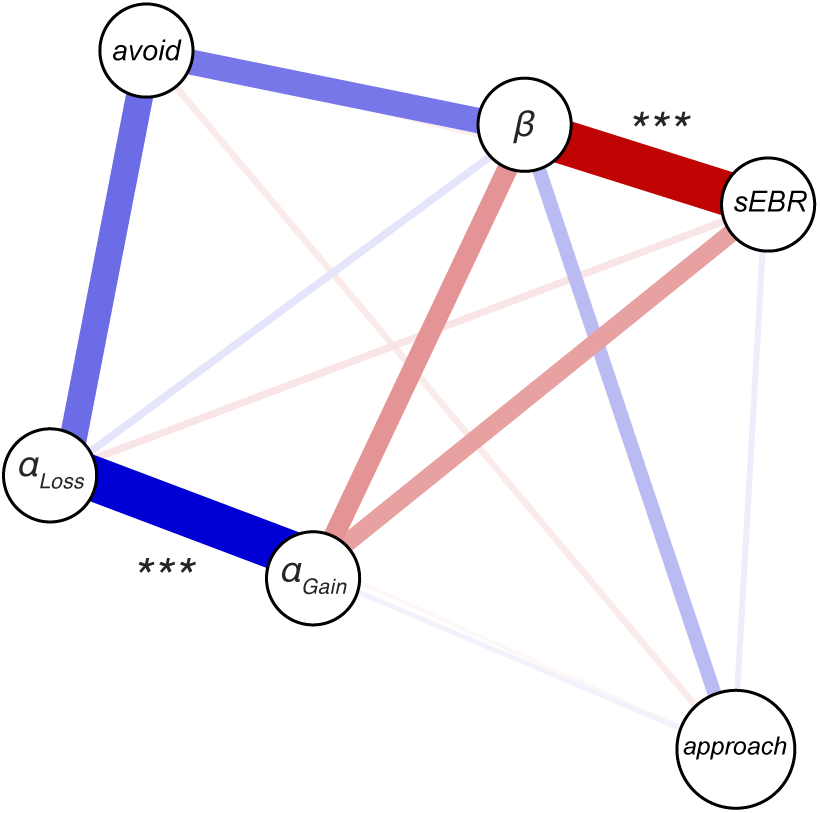
Network analysis. Graphical depiction of the partial correlation network of sEBR, approach A/avoid-B performance and the cognitive processes underlying learning. Variables of interest are represented as nodes. The estimated relations between variables are represented as edges, where the color of an edge (blue, red) indicates the direction of a relation (positive, negative) and the width of an edge indicates the strength of the observed relation. Edges are missing whenever the estimated relation between two nodes is zero. All nodes represent across-subject estimates. *β* = explore-exploit parameter; *α*_*Gain*_ = positive learning rate; *α*_*Loss*_ = negative learning rate; avoid = avoid-B accuracy, approach = approach-A accuracy.

## Discussion

The present study shows that performance on a probabilistic RL task is related to individual differences in sEBR. Our latent mixture modelling approach indicated that these learning differences were driven by a differential use of underlying cognitive processes, as we were able to distinguish individuals with low and high sEBR on the basis of their estimated learning rates and decision-making strategy. In addition, we found that sEBR uniquely predicted an individual’s explore-exploit tendency, thereby reflecting the sensitivity to value differences during a value-based choice. Specifically, choices of individuals with a lower sEBR were mostly determined by the relative value difference of presented options: they consistently exploited high valued options which resulted in better performance in the learning task. In contrast, individuals with a higher sEBR exhibited a more stochastic choice pattern with more frequent exploration of lower valued options, which resulted in lower learning phase performance. Our data suggest that variability in sEBR is related to an individual’s explore-exploit choice tendency during learning, with lower sEBR predicting stable, value-driven decisions, and higher sEBR predicting flexible, exploratory choices.

Our study investigated the link between sEBR and RL, but shows parallels with work in the field of cognitive flexibility, which is commonly described as the balance between maintaining stable task goals in the face of distraction versus flexible updating when the environment has changed^39^. In line with our finding that higher sEBR related to more explorative value-based choices, these studies have generally found that higher sEBR is associated with enhanced cognitive flexibility to support the detection of novel information in reversal learning^21^, working memory ^40^ and attentional set-shifting tasks^33,41-43^. For example, individuals with higher sEBR exhibited stronger tonic (or slow) pupil dilation during the detection of unexpected reward contingency reversals ^21^, suggestive of a stronger physical response to environmental change. Moreover, when the colour of a target switched to a novel one in an attentional set-shifting task, higher sEBR predicted improved task performance due to lower switching costs. In contrast, when the distractor colour became the target, higher sEBR predicted deteriorated performance due to higher switching costs as attention was drawn to the novel distractor colour^42^. As exploration or enhanced cognitive flexibility supports behaviour aimed at detecting novel information, this either improves or deteriorates performance depending on the environmental demands. In the learning phase of our task, participants experienced uncertainty due to the different reward probabilities of options, but not due to environmental change. Therefore, optimal task performance was achieved by making stable, exploitative choices for options with higher reward probabilities ^23^. This explains why individuals with a lower sEBR performed better in the certain AB and CD pairs, but not in the uncertain EF pair where more exploration was needed to discover the more often rewarded option.

Previous work that investigated sEBR in the context of probabilistic RL hypothesized that sEBR predicted how much individuals learned from the negative outcomes of their choices during prior learning^5^. This reasoning was based on the finding that sEBR correlated negatively with the ability to avoid the least rewarded option in a transfer phase that was administered after learning. As the relationship between sEBR and learning was not investigated directly, it remained unknown which cognitive process drove their observed effect. Both an exploitative decision-making strategy aimed at avoiding the least rewarded option and enhanced learning from negative feedback could explain the negative correlation between sEBR and avoidance of the least rewarded option. In the present study, we evaluated these alternative explanations directly by employing the Q-learning model that formalised learning and choice processes into learning rates and explore-exploit tendencies, respectively. We did not observe a relation between sEBR and negative learning rates (*α*_*Loss*_,), which indicated that sEBR did not relate to the magnitude by which participants learned from the negative outcomes of their choices as was hypothesized by Slagter et al., (2015). Consistently, sEBR was also unrelated to individuals’ learning rates after positive outcomes. Notable, while our analyses show that sEBR is not a reliable predictor of how participants learn from feedback, we find that it can be used as an index of one’s sensitivity to value differences during value-based choices, or explore-exploit tendency (*β*).

While we observed strong effects of sEBR during learning, effects on later value-based choices were rather weak or unreliable. Our network analysis - in which the unique relationship between two variables was estimated after controlling for the influence of all other variables - indicated that sEBR was unrelated to participants’ ability to avoid the least rewarded option. This finding is inconsistent with earlier work observing that sEBR did relate to the ability to avoid the least rewarded option^5^, or that it predicted the modulatory effect of dopaminergic drugs on approach and avoidance behaviours^6^. An important difference between these and our study, is that we evaluated both the effects of sEBR on learning as well as on later value-based choices, in one model. This analysis indicated that sEBR primarily related to an individual’s explore-exploit tendency during learning, that *in turn* related to the ability to avoid the least rewarded option in the transfer phase. Thus, individuals with a lower sEBR tended to exploit high valued outcomes, which especially improved learning in the AB pair and might be the reason that they avoided the least rewarded option B in the transfer phase. This could suggest that earlier observed effects of sEBR on approach and avoidance behaviours may be driven by individual variability in explore-exploit tendencies. Nevertheless, various studies have shown that seperate “Go” (approach-A) and “NoGo” neuronal populations (e.g. avoid-B) represent positive and negative action values that determine action selection^44-46^. Future studies that include dopaminergic manipulations combined with computational modelling to evaluate how sEBR relates to learning and later value-based choices might provide fruitful to answer this question.

Our observation that sEBR primarily reflects individual explore-exploit tendencies during learning could reconcile our work with the aforementioned studies^5,6^ as these and other studies^4,8,18^ have suggested that sEBR may reflect tonic, or baseline, striatal dopamine levels. Fluctuations in tonic dopamine levels have been observed to predominantly affect the expression, rather than learning, of motivated behaviour^47,48^, which agrees with our finding that sEBR uniquely predicted an individual’s explore-exploit tendency during a value-based choice. For example, several studies have shown that mice with chronically elevated tonic DA levels were more motivated to work for a food rewards, without showing improvements in Pavlovian or operant learning compared to wild type mice^49-51^. These experimental results are consistent with computational modelling studies that found that genetic or simulated differences in tonic DA levels uniquely correlated with explore-exploit tendencies, but not with learning rates^51-54^. Also in humans, some effects of dopaminergic medication on reward and punishment learning in PD patients can be explained by motivational differences at the time of choice, rather than by differences in feedback lea rning^55-57^. Together, these studies suggest that tonic DA levels impact the expression of motivated behaviour, or more specifically, explore-exploit tendencies. While our data preclude any conclusions about the biological mechanisms affecting sEBR, on the behavioural level our data agree with these studies linking sEBR to tonic striatal dopamine levels and individual variability in explore-exploit tendencies.

To conclude, sEBR predicted an individual’s tendency to explore or exploit during learning, thereby reflecting the sensitivity to value differences during a value-based choice. To our knowledge, this study is the first to associate sEBR to the underlying cognitive processes of learning, thereby providing a mechanistic understanding of the relation between sEBR, learning and the effects of learning on future value-based choices. We believe that using these methods advances our understanding of how sEBR relates to DA-dependent cognitive performance which could enable us to unify the diverse behavioural effects linked to sEBR, such as punishment or avoidance learning^5,6^, reversal learning^21^, as well as cognitive flexibility^33,41-43^. Together, our results indicate that sEBR can be used as an easy to measure behavioural index of individual explore-exploit tendencies during learning. Whether this is driven by fluctuations in tonic DA levels should be validated by other studies that directly measure or manipulate DA in a reinforcement learning task design.

## Methods

### Participants

The pupillometry data of the current data set was previously published^24^, but all sEBR data and analyses presented here are new. Forty-two healthy participants (10 males; mean age=24.9, range=18-34 years) with normal to corrected to normal vision participated participated in the experiment. Each participant was paid 16€ for two hours of participation and could earn an additional monetary bonus that depended on correct task performance (mean monetary bonus=10.2€, SD=1.8). The ethical committee of the Vrije Universiteit approved the study. All experimental protocols and methods described below were carried out in accordance with the guidelines and regulations of the Vrije Universiteit. Written informed consent was obtained from all participants. Four participants were excluded from analyses: one participant reported seeing more than three unique option pairs in the learning phase, and three participants had (almost) perfect choice accuracy in the learning phase, which complicated behavioural model fitting, leaving in total 38 participants for subsequent analyses.

### Blink rate recordings

Participants were seated in a dimly lit, silent room with their chin positioned on a chin rest, 60 cm away from the computer screen. An EyeLink 1000 Eye Tracker (SR Research) recorded at l000Hz seven minutes of spontaneous eye blinks from the continuously tracked eye data, which provides reliable sEBR estimates ^58^. Participants were kept naive about the sEBR measurements and were asked to maintain a normal gaze at a central fixation cross on the screen. All sEBR data was collected before 6 P.M., as sEBR is reported to be less stable during night time ^59^. Furthermore, participants were asked to sleep sufficiently the night before the experiment and to avoid the use of alcohol and other drugs of abuse.

### Task and procedure

After the blink rate recordings, participants performed a probabilistic RL task^60^ that consisted of a learning and a transfer phase. For an extended description of the task, stimuli and trial structure, we refer to^24^. Shortly, in the learning phase, participants completed 6 runs of 60 trials each (360 trials in total, 120 presentations of each option pair), with small breaks in-between runs. After each run, the earned number of points was displayed. At the end of the learning phase, the total number of earned points was converted into a monetary bonus.

Participants immediately proceeded to the transfer phase. In this phase, participants completed 5 runs of 60 trials each (300 trials in total, 20 presentations per option pair), with small breaks in-between runs. At the end of the transfer phase, choice accuracy across all trials was displayed and participants were fully debriefed about the sEBR measurements.

### Behavioural analyses

To assess how sEBR related to RL, we assigned each participant to the ‘low’ or ‘high’ sEBR group on the basis of a median split on across-subject sEBR values. We excluded two participants from analyses, as their sEBR fell exactly on the median, leaving 36 participants for subsequent analyses. A choice was regarded ‘correct’ when the option was chosen with the highest reward probability of each pair. Approach accuracy in the transfer phase was calculated as the percentage of trials in which the most rewarded option A was chosen when it was paired with another option. Avoidance accuracy was calculated as the percentage of trials in which the least rewarded option B was not chosen when it was paired with another option. In calculating approach and avoidance accuracy, the previous learning pairs (AB, CD, EF) were excluded to account for repetition effects.

### Q-leaming model

To investigate how sEBR related to the cognitive processes underlying RL, we applied a Q-learning model^1,61^ to each participant’s sequence of choices in the learning phase. During Q-learning, individuals update their value belief, or “Q-value”: of the recently chosen option by learning from feedback that resulted in an unexpected outcome. All Q-values were initialised at 0.5. Learning is captured by the reward prediction error (RPE) and can be formally described by a delta rule:

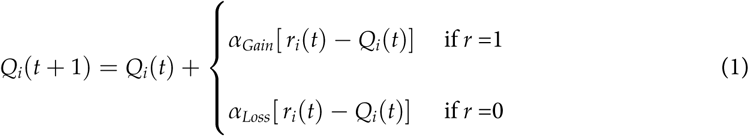

where parameters 0 ≤ *α*_*Gain*_, *α*_*Loss*_ ≤ 1 represent positive and negative learning rates, that independently regulate the impact of recent positive and negative feedback on current value beliefs. A relatively high learning rate indicates more sensitivity to recent feedback, whereas a relatively low learning rate indicates a stronger focus on the integration of feedback over multiple trials^27^. A choice between two presented stimuli on the next trial was described by a “softmax” choice rule:

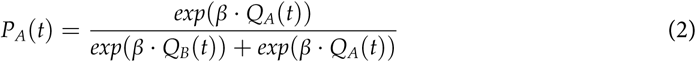

Here, 0 ≤ *β* ≤ 100, or the explore-exploit parameter, describes an individual’s sensitivity to value differences between presented stimuli, where a higher *β* value indicate greater sensitivity to smaller value differences, hence, exploitative choices for high reward options **(Figure 1b**).

### Bayesian hierarchical implementation of the Q-leaming model

We implemented the Q-learning model in a hierarchical Bayesian framework **(Supplementary Figure 1**)^22-24,62^, in which group-level and individual-level parameter distributions are simultaneously fit that mutually constrain each other. This approach results in greater statistical power and more stable parameter estimation compared to procedures using individual-level maximum likelihood^63,64^. To examine the cognitive processes underlying learning for low and high sEBR groups, we fit separate group-level parameter distributions of positive and negative learning rates (*α*_*Loss*_, *α*_*Gain*_) and explore-exploit tendencies(*β*). For an extended description of the applied Bayesian hierarchical model, we refer to^24^.

### Bayesian latent mixture modelling

We performed Bayesian latent mixture modelling on participants’ choice data in the learning phase to assess whether an individual’s sEBR could be predicted on the basis of the estimated cognitive processes (*α*_*Loss*_, *α*_*Gain*_ and *β*) underlying learning **(Figure 1c)**^31,65^.We evaluated all participants in one dataset and discarded information about their measured sEBR. Importantly, we still assumed that each participant belonged to either of the two sEBR groups, but that their group membership had to be determined. Thus, the goal of this analysis was to investigate whether a participan t’s sEBR group membership can be inferred from the estimated cognitive processes alone.

To estimate a participant’s group membership, we used a binary indicator variable *x*_*i*_, where *x*_*i*_ = 0 and *x*_*i*_ = 1 indicates that participant *i* belongs to the low and high sEBR group, respectively. For each participant, the posterior mean of the *x*_*i*_ variables reflected the probability to be classified into the high sEBR group. Following Steingroever et al. (2017), we used informative priors to inform the group membership indicator variable during model fitting. These priors were derived from the previous Bayesian hierarchical parameter analyses, and approximated the group-level posterior parameter distributions (*α*_*Gain*_, *α*_*Loss*_ and *β*) for the low and high sEBR groups. Specifically, for each group probit transformed individual-level parameters were drawn from a group-level normal distributions *z*′ ∼ 𝒩 (*μ*_*z*_, *σ*_*z*_). These normal prior distributions were characterised by each group’s mean and standard deviation that we derived from the posterior distributions of our previous model fits. Thus, the group-level posterior parameter distributions of low and high sEBR groups were used as informative prior distributions for the latent mixture modelling analysis. It is important to note that the mixture model was at all times blind about each participant’s sEBR group membership. This was predicted by modelling each participant’s choice data and evaluation against the group-level priors. As we used the behavioural data both to construct the prior distributions and to fit the latent mixture model, we cannot make inferences about the model parameters^31^. However, this analysis provides a way to investigate whether a participant’s sEBR group membership can be inferred on the basis of the cognitive processes that drive RL.

### Model estimation

Our model-based analyses were implemented in PyStan mc-stan.org and fit to all trials of the learning phase that fell within the correct response time window 150ms ≤ RT ≤ 3500ms. We ran four Markov Chain Monte Carlo (MCMC) chains for both the Bayesian hierarchical parameter estimation and latent mixture model, of which we collected 5000 and 9000 samples each (after discarding the first 1000 samples of each chain for burn-in). Visual inspection of the chains suggested the model converged. This was validated by the Rhat statistic^63^, a convergence diagnostic that compares between and within chain variability, as all Rhats were <1.05. Simulations displayed in **Supplementary Figure 2** were similar to observed choice behavior for both the low and high sEBR groups.

### Multiple regression analyses

We performed frequentist and Bayesian multiple regression analyses in JASP jasp-stats.org to quantify the relative influence of each model parameter (*α*_*Gain*_, *α*_*Loss*_ and *β*) on 1) individual variability in sEBR and 2) approach/avoidance behaviour in the transfer phase. For all Bayesian multiple regression analyses we used the default priors from JASP. Bayesian multiple regression analyses in JASP follow a model comparison approach, in which the influence of each parameter and combinations thereof are evaluated step by step. Resulting Bayes Factors (BF) are interpreted as the odds supporting one model over another. BF-values between 3-10 indicate substantial support for the alternative model over the null model that a regressor’s true value is zero, whereas BF-values > 10 indicate strong support that the alternative model is favoured over the null model^37^. For all analyses, we selected the modes of the individual posterior parameter distributions of all participants. These variables were log-transformed and normalised prior to analysis to account for parameter skewness and scaling effects.

## Network analysis

We performed a network analysis in JASP, in which the relation between any two variables in the network is estimated directly while accounting for the influence of all other variables in the network. Thus, the analysis reflects the unique relationship between two variables that cannot be explained by or result from other factors. We estimated a partial correlations network to capture the unique relationships between 1) sEBR, 2) the cognitive processes driving learning (*α*_*Gain*_, *α*_*Loss*_, and *β*), and 3) approach-A and avoid-B choices in the subsequent transfer phase.

## Acknowledgements

The authors would like to thank Lisa Roodermond and Lynn van den Berg for their assistance in the data collection of this study. This research is funded by the ERC advanced grant [ERC-2012-AdG-323413] to JT.

## Author Contributions Statement

SJ and JS conceptualised research ideas and designed analyses. SJ contributed novel analytical methods. JS collected and analysed the data. JS and SJ wrote the original draft. JS prepared the figures. JS, SJ and JT edited and reviewed the manuscript. SJ supervised the project. JT funded the project.

## Data availability

The OSF DOI link to the raw data and analysis scripts is: 10.17605/OSF.IO/4PQ9C

