## Supplementary Materials for "Spontaneous eye blink rate predicts individual differences in exploration and exploitation during reinforcement learning"

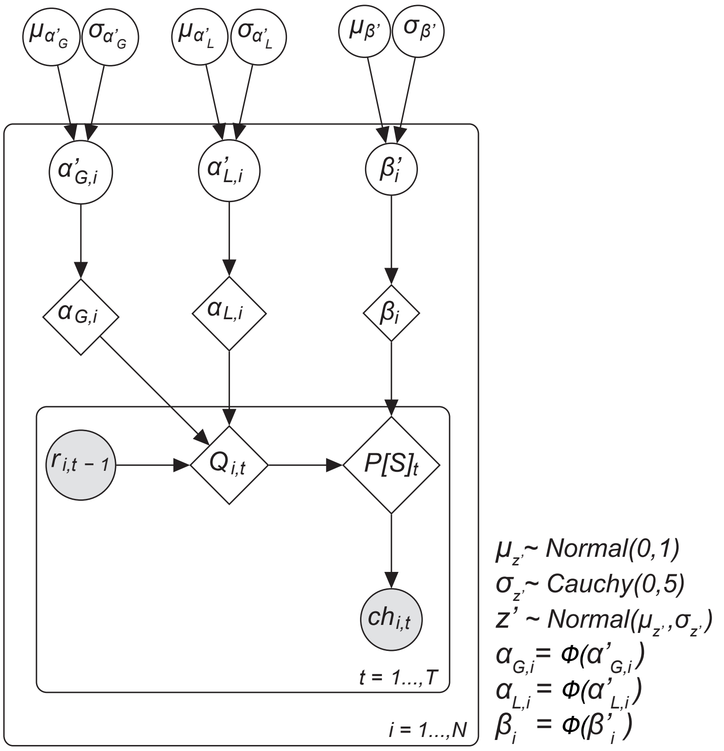


***Supplementary Figure 1.*** **Graphical representation of the hierarchical Bayesian Q-learning model.** The inner plane represents within-subject trial-by-trial RL behaviour. Variables $r_i\,(t\text{-}1)$ (outcome for participant *i* on trial *t-1*) and $ch_i(t)$ (choice of participant *i* on trial *t*) were obtained from the behavioural data. The outer plane represents per-participant parameter estimates $\alpha_{Gi}$ ($\alpha_{Gain}$ participant *i*), $\alpha_{Li}$ ($\alpha_{Loss}$ participant *i*) and $\beta_i$ ($\beta$ participant *i*) that were fit separately for participants in the low and high sEBR group. Per-participant parameter estimates were modelled using a probit transform $z'_i$ ($\alpha'_{Gi}$, $\alpha'_{Li}$, $\beta'_i$). $z'_i$ were drawn from group-level normal distributions with mean $\mu_{z\prime}$ and standard deviation $\sigma_{z\prime}$. The outermost layer represents group-level mean and standard deviations of the Q-learning model parameters. A normal prior was assigned to all group-level means, $\mu_{z\prime}$~*N(0,1),* and a half-Cauchy prior to all group-level standard deviations, $\sigma_{z\prime}$~*Cauchy(0,5).* A weakly informative prior such as this is recommended in small sample sizes to reduce the influence of the priors on posterior distributions[66]. Shaded variables are obtained from the behavioural data and used to fit the model. Diamond shaped nodes are deterministic, as they are derived from the model fit. Circular unshaded nodes indicate continuous variables. Arrows indicate dependencies between variables. *Φ()* represents the probit transform.

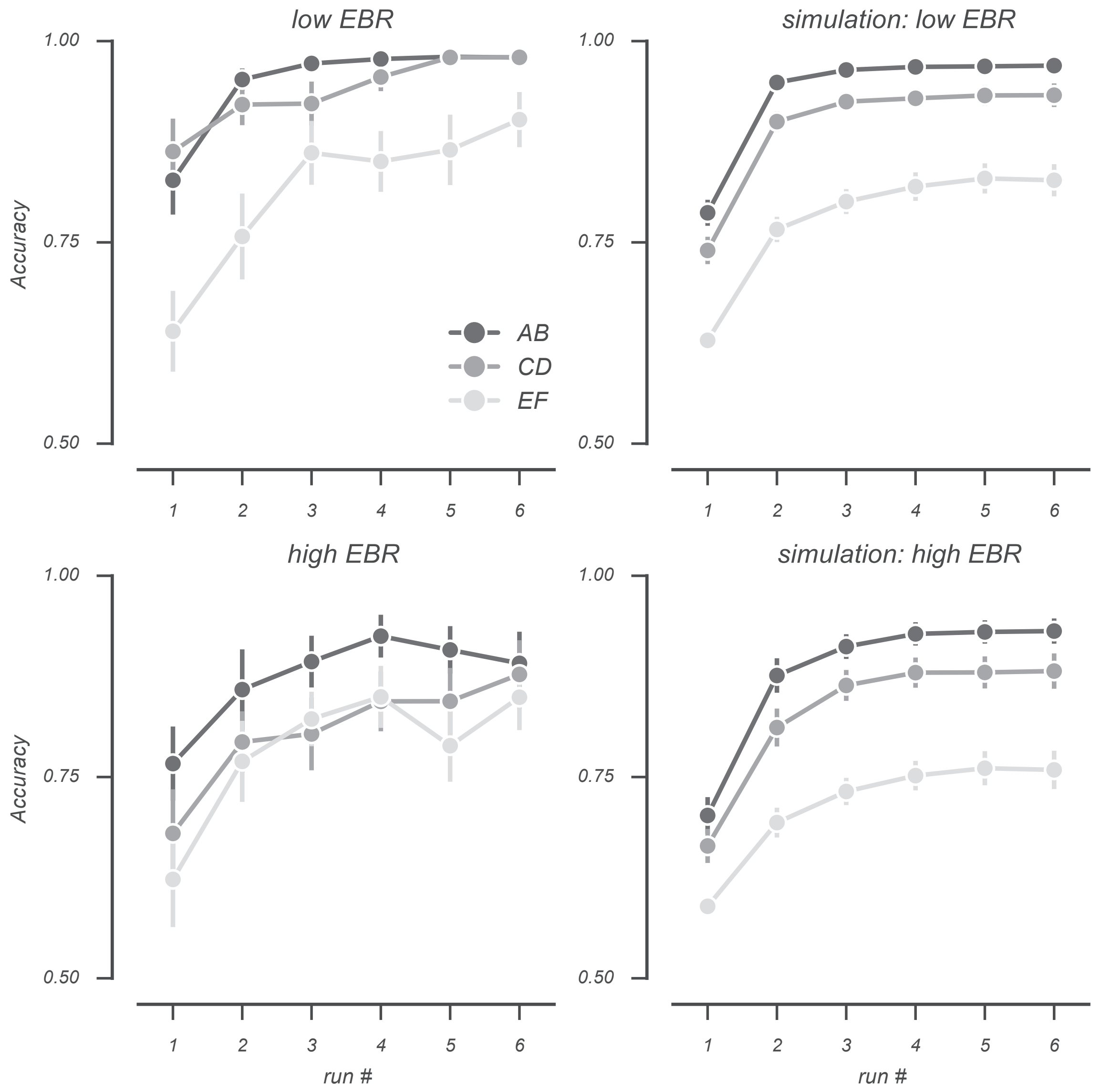


***Supplementary Figure 2. Real and simulated learning phase choice accuracy for low and high sEBR groups***. Choices were simulated by drawing 100 samples from each participant's posterior parameter distributions. Real and simulated choice accuracy was split by run number and option pair and averaged across participants. All simulations approached observed choice behavior, with the exception that EF choice accuracy was underestimated by the model for individuals with a high sEBR.

**Model Comparison**

| Models | P(M) | P(M ǀ data) | $BF_M$ | $BF_{10}$ | $R^2$ |
|---|---|---|---|---|---|
| Null model | 0.125 | 0.007 | 0.052 | 1.000 | 0.000 |
| $\beta$ | 0.125 | 0.353 | 3.814 | 47.859 | 0.297 |
| $\beta + \alpha_{Gain}$ | 0.125 | 0.314 | 3.205 | 42.618 | 0.349 |
| $\beta + \alpha_{Loss}$ | 0.125 | 0.196 | 1.710 | 26.634 | 0.327 |
| $\beta + \alpha_{Gain} + \alpha_{Loss}$ | 0.125 | 0.116 | 0.917 | 15.722 | 0.352 |
| $\alpha_{Loss}$ | 0.125 | 0.006 | 0.045 | 0.868 | 0.071 |
| $\alpha_{Gain}$ | 0.125 | 0.004 | 0.031 | 0.598 | 0.045 |
| $\alpha_{Gain} + \alpha_{Loss}$ | 0.125 | 0.003 | 0.020 | 0.389 | 0.077 |

***Supplementary Table 1. Bayesian linear regression analysis of Q-learning model parameter modes on sEBR.*** Compared to the null model, the data provide strong evidence in favour of the model in which the β-parameter explains individual variability in sEBR.

**Model Comparison**

| Models | P(M) | P(M ǀ data) | $BF_M$ | $BF_{10}$ | $R^2$ |
|---|---|---|---|---|---|
| Null model | 0.125 | 0.048 | 0.350 | 1.000 | 0.000 |
| $\beta + \alpha_{Loss}$ | 0.125 | 0.338 | 3.574 | 7.108 | 0.260 |
| $\alpha_{Loss}$ | 0.125 | 0.168 | 1.411 | 3.527 | 0.159 |
| $\beta + \alpha_{Gain} + \alpha_{Loss}$ | 0.125 | 0.134 | 1.079 | 2.809 | 0.260 |
| $\beta$ | 0.125 | 0.133 | 1.069 | 2.786 | 0.145 |
| $\beta + \alpha_{Gain}$ | 0.125 | 0.092 | 0.713 | 1.944 | 0.185 |
| $\alpha_{Gain} + \alpha_{Loss}$ | 0.125 | 0.064 | 0.475 | 1.336 | 0.161 |
| $\alpha_{Gain}$ | 0.125 | 0.025 | 0.177 | 0.518 | 0.035 |

***Supplementary Table 2. Bayesian linear regression analysis of Q-learning model parameter modes on avoidance accuracy in the transfer phase.*** Compared to the null model, the data provide moderate evidence in favour of the model in which both the β-parameter and $\alpha_{Loss}$-parameter explain individual variability in avoidance behavior in the transfer phase.

**Weights matrix**

| Variable | Network | | | | | |
|---|---|---|---|---|---|---|
| | $\alpha_{Gain}$ | $\alpha_{Loss}$ | approach | avoid | β | sEBR |
| $\alpha_{Gain}$ | 0.000 | 0.522* | -0.016 | -4.340e -4 | -0.220 | -0.192 |
| $\alpha_{Loss}$ | 0.522* | 0.000 | 0.026 | 0.302 | 0.055 | -0.053 |
| approach | -0.016 | 0.026 | 0.000 | -0.043 | 0.139 | 0.037 |
| avoid | -4.340e -4 | 0.302 | -0.043 | 0.000 | 0.278 | -0.034 |
| β | -0.220 | 0.055 | 0.139 | 0.278 | 0.000 | -0.515* |
| sEBR | -0.192 | -0.053 | 0.037 | -0.034 | -0.515* | 0.000 |

***Supplementary Table 3. Partial correlation weights matrix of all network variables.*** Asterisks indicate significant partial correlations between variables in the network.

### *Supplementary references*

66. Ahn, W-Y., Haines, N. & Zhang, L. Revealing neurocomputational mechanisms of reinforcement learning and decision-making with the hBayesDM Package. *Computational Psychiatry,* ***1****, 24-57 (2017).*
